# NRF1 and NRF3 complementarily maintain a basal proteasome activity in cancer cells through CPEB3-mediated translational repression

**DOI:** 10.1101/2020.01.10.902718

**Authors:** Tsuyoshi Waku, Hiroyuki Katayama, Miyako Hiraoka, Atsushi Hatanaka, Nanami Nakamura, Yuya Tanaka, Natsuko Tamura, Akira Watanabe, Akira Kobayashi

## Abstract

Proteasomes are protease complexes essential for cellular homeostasis, and their activity is crucial for cancer cell growth. However, the mechanism of how proteasome activity is maintained in cancer cells has remained unclear. The CNC family transcription factor NRF1 induces the expression of almost all proteasome-related genes under proteasome inhibition. *NRF1* and its phylogenetically closest homolog *NRF3* are both highly expressed in several types of cancers, such as colorectal cancer. Herein, we demonstrate that NRF1 and NRF3 complementarily maintain basal proteasome activity in cancer cells. A double knockdown of *NRF1* and *NRF3* impaired the basal proteasome activity in cancer cells and the cancer cell resistance to a proteasome inhibitor anticancer drug bortezomib by significantly reducing basal expression of seven proteasome-related genes, including *PSMB3, PSMB7, PSMC2, PSMD3, PSMG2, PSMG3*, and *POMP*. Interestingly, the molecular basis behind these cellular consequences was that NRF3 repressed NRF1 translation by the gene induction of translational regulator CPEB3, which binds to *NRF1*-3′UTR and decreases polysome formation on *NRF1* mRNA. Consistent results were obtained from clinical analysis, wherein patients with cancer having higher *CPEB3*/*NRF3*-expressing tumors exhibit poor prognosis. These results provide the novel regulatory mechanism of basal proteasome activity in cancer cells through an NRF3-CPEB3-NRF1 translational repression axis.

## Introduction

Cancer cell survival and growth is dependent on the activity of a proteasome, which is a large protease complex that catalyzes the precise and rapid degradation of proteins involved in antitumorigenic events, such as cell-cycle arrest and apoptosis induction^1,2^. One of the proteasome regulators in mammals is the cap’n’collar (CNC) family transcription factor, NRF1 (NFE2L1). NRF1 induces the expression of almost all proteasome-related genes under proteasome inhibition^3–6^, and the brain-specific knockout actually impairs proteasome function and causes neurodegeneration in mice^7^. However, the transcriptional regulation of proteasome activity in cancer cells has remained unclear.

Phylogenetically, NRF3 (NFE2L3) has been identified as the closest homolog of NRF1 in the CNC family^8^. Recently, *NRF3* gene amplification has been reported in patients with colorectal cancer^9^. In the Gene Expression Profiling Interactive Analysis (GEPIA) web server^10^, the *NRF3* gene is highly expressed in many types of tumors, such as testicular germ cell tumors, pancreatic adenocarcinoma, colon adenocarcinoma (COAD), and rectum adenocarcinoma (READ), whereas the *NRF1* gene is highly expressed in almost all types of normal and tumor tissues. These insights imply the functional correlation between NRF1 and NRF3 with respect to the maintenance of a basal proteasome activity in cancer cells.

In this study, we show that both NRF1 and NRF3 are required to maintain a basal proteasome activity in cancer cells through the induction of several proteasome-related genes, including *PSMB3, PSMB7, PSMC2, PSMD3, PSMG2, PSMG3*, and *POMP*. Interestingly, NRF3 represses NRF1 translation by inhibiting polysome formation on *NRF1* mRNA. We identify a translational regulator *CPEB3* (*cytoplasmic polyadenylation element-binding protein 3*) as an NRF3-target gene that is involved in the repression of NRF1 translation. A functional CPEB recognition motif is also identified in the *NRF1*-3′UTR. Furthermore, we validate that CPEB3 is a key factor for not only the complementary maintenance of a basal proteasome activity by NRF1 and NRF3 but also the poor prognosis of colorectal cancer patients with tumors highly expressing *NRF3*, but not *NRF1*. In conclusion, we demonstrate the novel regulatory mechanism of basal proteasome activity in cancer cells where NRF1 and NRF3 complementarily, but not simultaneously, maintain the basal expression of proteasome-related genes through CPEB3-mediated translational repression.

## Results

### Both NRF1 and NRF3 are required to maintain a basal proteasome activity in cancer cells

Initially, we investigated the biological relevance of NRF1 and NRF3 for proteasome activity at the basal level in living cancer cells. Using human colorectal carcinoma HCT116 cells, we generated cells which stably expressed the ZsProSensor-1 fusion protein, a proteasome-sensitive fluorescent reporter (Supplementary Fig. 1A). Once the proteasome in this stable cell line was inhibited by proteasome inhibitor MG-132, green fluorescence derived from the reporter protein was detected using a flow cytometer (Supplementary Fig. 1B). We found that a double knockdown of *NRF1* and *NRF3* significantly decreased basal proteasome activity in living cancer cells (Fig. 1A). Consistent results were obtained by *in vitro* proteasome activity assays (Supplementary Fig. 1C). The double knockdown also impaired the cancer cell resistance to a proteasome inhibitor bortezomib (BTZ), which is clinically used as an anticancer drug^11,12^ (Fig. 1B).

**Figure 1.**
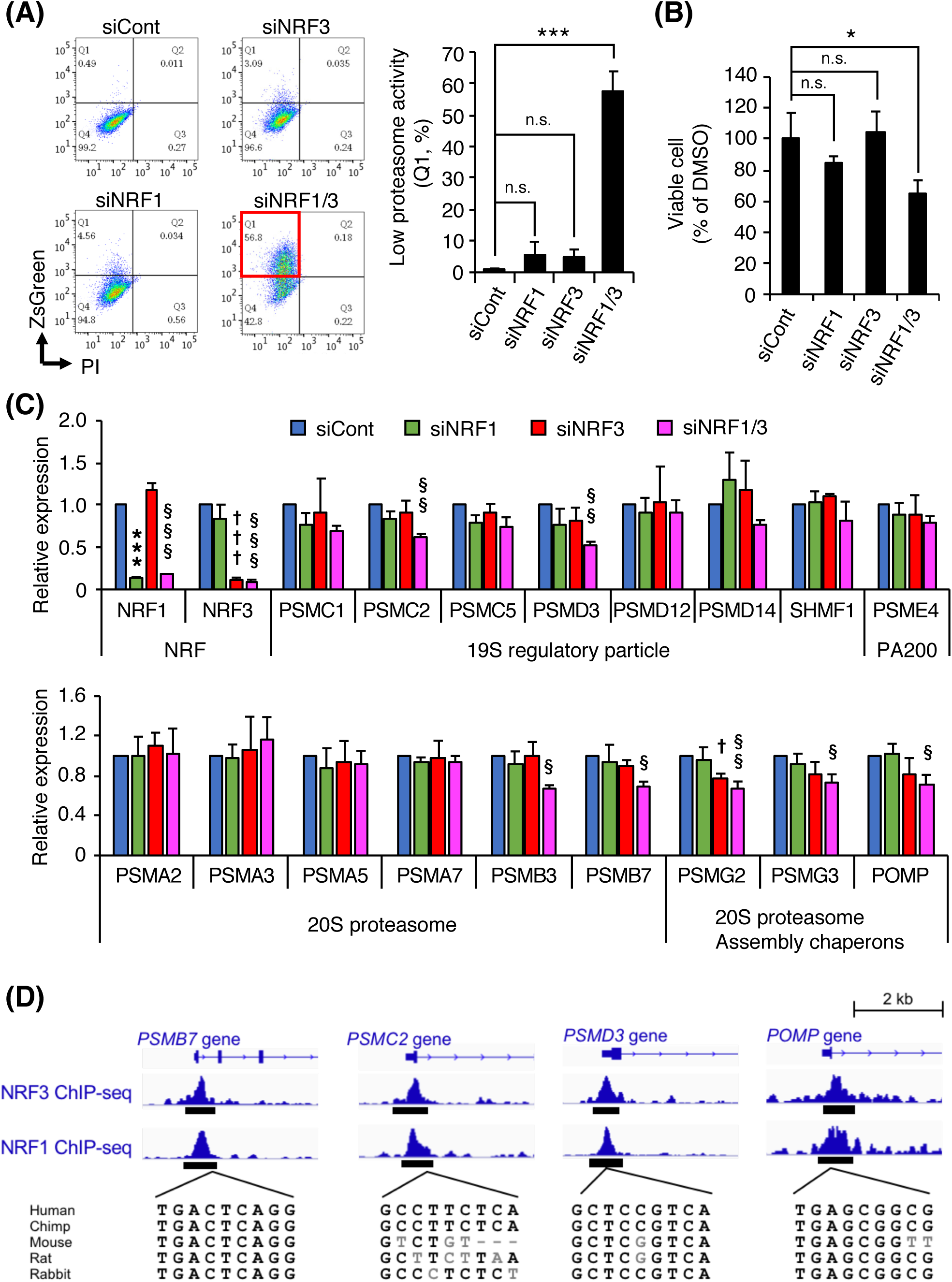
NRF1 and NRF3 complementarily regulate proteasome activity and proteasome subunit gene expression at a basal level. (**A**) Impact of *NRF* knockdown on a basal proteasome activity. At 24 hours after siRNA transfection into ZsPS cells, the fluorescence intensity derived from a ZsProSensor-1 reporter was measured using flow cytometry. The cell populations in Q1 enclosed by a red line are those with “low proteasome activity.” siNRF1/3 is a double knockdown of *NRF1* and *NRF3*. Control siRNA (siCont) was used as a control. \*\*\**p* < 0.005; n.s., not significant (*n* = 3, mean ± SD, ANOVA followed by Tukey’s test). (**B**) Impact of *NRF* knockdown on the BTZ resistance. At 24 hours after siRNA transfection to HCT116 cells, the cells were treated with 5 nM BTZ and further incubated for 48 hours. Then, the cells were subjected to cell viability assay using trypan blue staining. siNRF1/3 represents a double knockdown of *NRF1* and *NRF3*. Control siRNA (siCont) and DMSO were used as controls. Cell viability was determined by the number of living cells with BTZ treatment normalized by that with DMSO. \**p* < 0.05; n.s., not significant (*n* = 3, mean ± SD, ANOVA followed by Tukey’s test) (**C**) Impact of *NRF* knockdown on mRNA levels of 17 common core genes with a “Yes” value in the core enrichment in both *NRF1* and *NRF3* knockdown cells (Supplementary Table 2). At 48 h after siRNA transfection into HCT116 cells, the cells were analyzed by RT-qPCR. siNRF1/3 represents a double knockdown of *NRF1* and *NRF3*. Control siRNA (siCont) was used as a control. mRNA levels of each proteasome subunit were normalized according to levels of *β-actin* mRNA. \**p* < 0.05; \*\**p* < 0.01; \*\*\**p* < 0.005 (*n* = 3, mean ± SD, ANOVA followed by Tukey’s test, *siCont vs. siNRF1; †siCont vs. siNRF3; §siCont vs. siNRF1/3) (**D**) ChIP peaks of NRF1 and NRF3 in the promoters of proteasome-related genes. ChIP sequencing of endogenous NRF1 or exogenous NRF3 was performed using wild-type HCT116 cells or *NRF3*-overexpression H1299 cells treated with 1 μM MG-132 for 24 hours.

Next, we investigated the impact of NRF1 and NRF3 on the expression of proteasome-related genes. To address this issue, we performed DNA microarray analysis using *NRF1*- or *NRF3*-knockdown HCT116 cells, and found 42 proteasome-related genes with a decrease in expression of less than 1.4-fold (Supplementary Table 1). Gene Set Enrichment Analysis (GSEA) using these array datasets showed reduced expression of 17 common core genes in both *NRF1* and *NRF3* knockdown cells (Supplementary Fig. 1D and E, Supplementary Table 2). Therefore, using RT-qPCR we showed that the mRNA levels of *PSMB3, PSMB7, PSMC2, PSMD3, PSMG2, PSMG3*, and *POMP* were significantly decreased by the double knockdown of *NRF1* and *NRF3* (Fig. 1C). These results indicate that both NRF1 and NRF3 are required to maintain the basal expression of several proteasome-related genes.

NRF1 induces proteasome-related genes by directly binding antioxidant response elements (ARE) in their gene promoters^3–6^. NRF3 also binds to ARE sequence *in vitro*^8^, although it has not been reported whether NRF3 binds to the ARE in the promoters of proteasome-related genes in cells. To address this issue, we analyzed our chromatin immunoprecipitation (ChIP) sequencing datasets in the presence of proteasome inhibitor MG-132, which stabilizes both NRF1 and NRF3 proteins (manuscript in preparation) and found the positive ChIP peak of NRF1 and NRF3 proteins on the promoters of *PSMB7, PSMC2, PSMD3*, and *POMP* genes (Fig. 1D), suggesting that NRF3 as well as NRF1 directly induces the basal expression of several proteasome-related genes.

### NRF3 represses NRF1 translation by inhibiting polysome formation on *NRF1* mRNA

To clarify the molecular mechanism behind the maintenance of a basal proteasome activity in cancer cells by both NRF1 and NRF3, we then investigated the relationship between NRF1 and NRF3 expression in HCT116 cells. Interestingly, NRF1 protein levels were increased by *NRF3* knockdown, while NRF3 protein levels were not changed by *NRF1* knockdown (Fig. 2A). Similar results were obtained in other cancer cell lines, including T98G (human glioblastoma multiforme), and MCF-7 (human breast cancer) (Supplementary Fig. 2A). The levels of *NRF1* mRNA were unchanged by *NRF3* knockdown (Fig. 2B, Supplementary Fig. 2B). We also obtained consistent results in cells in which NRF3 was overexpressed (Supplementary Fig. 2C and D). These results indicate that NRF3 decreases NRF1 protein levels without altering *NRF1* gene expression, suggesting the possible effect of NRF3 on the protein stability or translation of NRF1. To investigate the former effect, we performed a cycloheximide (CHX) chase experiment and found that *NRF3* knockdown did not stabilize NRF1 proteins (Fig. 2C). Meanwhile, we investigated the latter effect by a polysome profiling analysis and showed that *NRF3* knockdown increased the amount of *NRF1* mRNA in polysomes between fractions #11 and #20 (Fig. 2D), although rRNA distribution and global protein synthesis remained unchanged (Supplementary Fig. 2E and F). These results indicate that NRF3 decreases polysome formation on *NRF1* mRNA, thereby repressing NRF1 translation.

**Figure 2.**
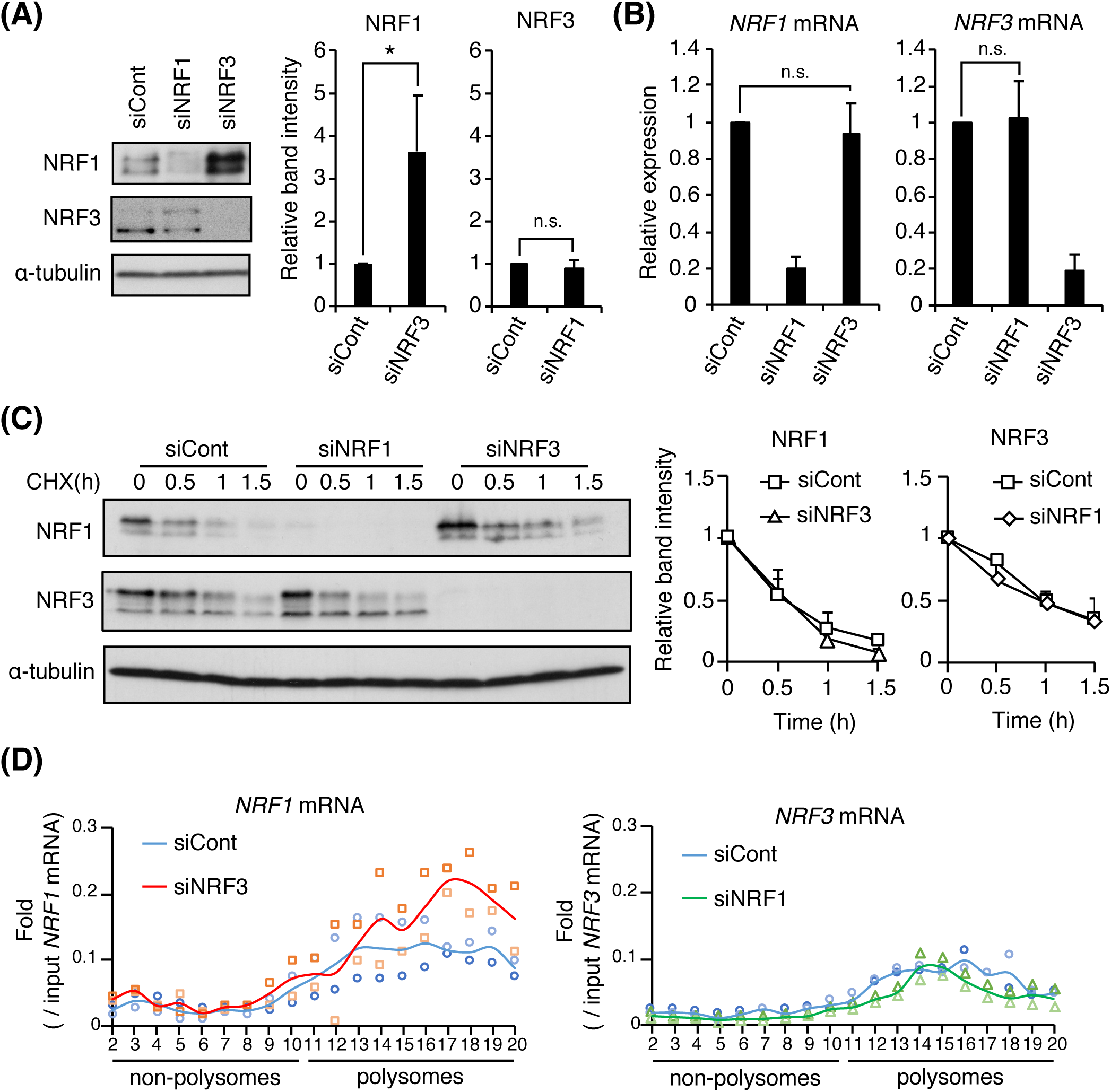
NRF3 represses NRF1 translation by inhibiting polysome formation on *NRF1* mRNA. (**A** and **B**) Impact of *NRF* knockdown on protein and mRNA levels of another NRF in HCT116 (colorectal carcinoma) cells. At 48 hours after siRNA transfection, the cells were analyzed by immunoblotting (**A**) and RT-qPCR (**B**). Protein or mRNA levels of each NRF were normalized with reference to α-tubulin protein or *β-actin* mRNA, respectively. Control siRNA (siCont) was used as a control. \**p* < 0.05; n.s. not significant (*n* = 3, mean ± SD, t-test in **A**, ANOVA followed by Tukey’s test in **B**) (**C**) Impact of *NRF* knockdown on protein degradation of another NRF. At 48 hours after siRNA transfection, the cells were treated with CHX, and analyzed by immunoblotting at the times indicated. Protein levels were normalized by α-tubulin. Control siRNA (siCont) was used as a control. (*n* = 3, mean ± SD) (**D**) Distribution of *NRF* mRNA in HCT116 transfected with indicated siRNA (*n* = 2). At 48 hours after siRNA transfection, the cells were analyzed by sucrose gradient centrifugation followed by RT-qPCR. mRNA levels in each fraction were normalized against those in the input. Mean and individual values are represented as a line and mark, respectively. Control siRNA (siCont) was used as a control.

### NRF3 directly induces the gene expression of translational regulator *CPEB3*

To identify an NRF3-target gene which represses NRF1 translation, we performed DNA microarray analysis using *NRF3*-knockdown HCT116 cells and *NRF3*-overexpression H1299 (human non-small cell lung cancer) cells and identified 146 genes whose expression was positively associated with *NRF3* gene expression (Supplementary Fig. 3A). Among these genes, we found *CPEB3* to be a candidate gene related to translational regulation, using gene ontology analysis (Supplementary Fig. 3B) and validated this result using RT-qPCR (Fig. 3A). We then confirmed that NRF3 proteins bound to one of two ARE regions within the *CPEB3* promoter (Fig. 3B and C), indicating that NRF3 directly induces *CPEB3* gene expression. We investigated whether CPEB3 is related to the NRF3-mediated regulation of NRF1 translation and found that *CPEB3* knockdown increased NRF1 protein levels and the amount of *NRF1* mRNA in polysomes, while rRNA distribution and global protein synthesis remained unchanged (Fig. 3D and E, Supplementary Fig. 3C–E). We further confirmed that NRF1 protein levels were decreased by *CPEB3* overexpression (Fig. 3F). These results clearly demonstrate that NRF3 specifically represses NRF1 translation in a CPEB3-dependent manner. We also found evidence of *CPEB3* gene induction by NRF1 (Supplementary Fig. 3F), suggesting the presence of negative feedback regulation of NRF1 through CPEB3-mediated translation repression. We discuss the biological implications of this finding in the Discussion section.

**Figure 3.**
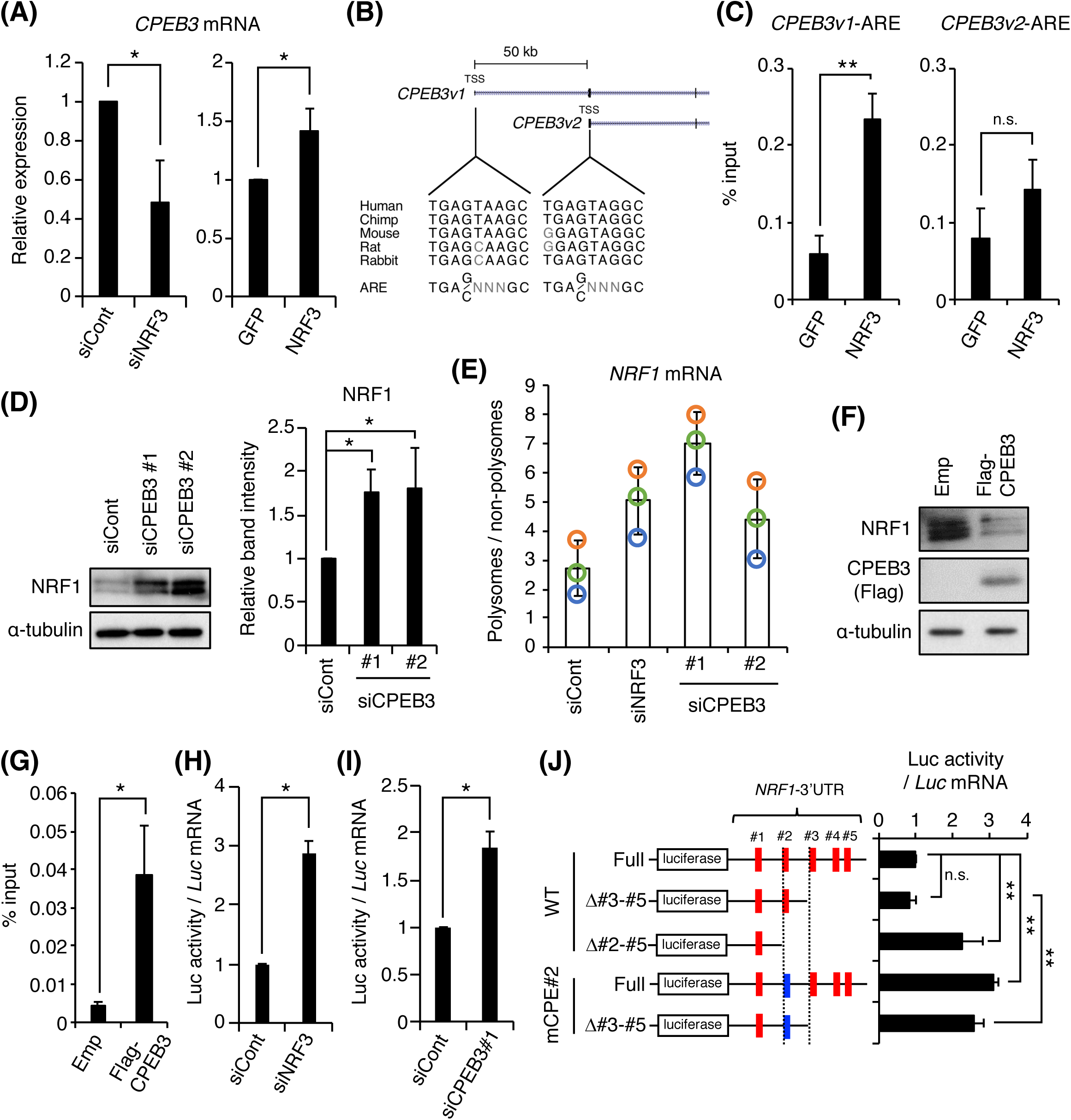
CPEB3 is an NRF3-target gene that negatively regulates NRF1 translation. (**A**) Impact of *NRF3* knockdown on mRNA levels of *CPEB3* in HCT116 cells (left pane) and *NRF3*-overexpression H1299 cells (right panel). Control siRNA (siCont) or *GFP*-overexpression H1299 cells were used as controls. mRNA levels of *CPEB3* were normalized according to *β-actin* mRNA. \**p* < 0.05 (*n* = 3, mean ± SD, t-test) (**B** and **C**) The recruitment of NRF3 on *CPEB3* promoters in *NRF3*-overexpression H1299 cells. *GFP*-overexpression H1299 cells were used as controls. In (**B**), the genome locus of the *CPEB3* promoter and multiple sequences of two candidate AREs in indicated species are shown. TSS; Transcription start site. \*\**p* < 0.01; n.s., not significant (*n* = 3, mean ± SD, t-test) (**D**) Impact of *CPEB3* knockdown on NRF1 protein levels. Each siRNA was transfected into HCT116 cells. After 48 hours, the cells were analyzed by immunoblotting. Representative images of immunoblotting are shown in the left panel, and the protein levels were normalized by α-tubulin in the right panel. Control siRNA (siCont) was used as a control. \**p* < 0.05 (*n* = 3, mean ± SD, ANOVA followed by Tukey’s test) (**E**) Impact of *CPEB3* knockdown on the amount of *NRF1* mRNA on polysomes. At 48 hours after siRNA transfection, the cells were subjected to sucrose gradient centrifugation. The fractions from #2 to #8 and from #11 to #19 shown in Supplementary Fig. 3D were collected as the non-polysomal and polysomal fractions, respectively. Each fraction was analyzed by RT-qPCR. mRNA levels of *NRF1* in each fraction were normalized against those in the input. *NRF1* mRNA levels in the polysomal fractions were then divided by *NRF1* mRNA levels in the non-polysomal fractions. Control siRNA (siCont) and siNRF3 were used as controls. Mean ± SD and three independent values are represented as bars and indicated colored marks, respectively (*n* = 3). (**F**) Impact of *CPEB3* overexpression on NRF1 protein levels. At 48 hours after 3×Flag-CPEB3 plasmid transfection into HCT116 cells, the cells were analyzed by immunoblotting. Empty vector (Emp) was used as a control. (**G**) Interactions between CPEB3 protein and *NRF1* mRNA. At 24 hours after 3×Flag-CPEB3 plasmid transfection, cells were analyzed by RIP assay followed by RT-qPCR. RIPed mRNA levels of *NRF1* were normalized against the input values. Empty vector (Emp) was used as a control. \**p* < 0.05 (*n* = 3, mean ± SD, t-test) (**H** and **I**) Impact of *NRF3* or *CPEB3* knockdown on NRF1 translation *via* its 3′UTR. At 24 hours after the transfection of siNRF3 (**H**) or siCPEB3 (**I**), a luciferase reporter vector fused with *NRF1*-3′UTR was transfected and culture for 24 hours. Control siRNA (siCont) was used as a control. Luciferase activity was normalized by mRNA levels of a *luciferase* gene. \**p* < 0.05 (*n* = 3, mean ± SD, t-tests) (**J**) Impact of *NRF1*-3′UTR deletion or mutation on translation. The luciferase reporter vector fused with the indicated *NRF1*-3′UTR was transfected into HCT116 cells, and the cells were analyzed after 24 hours. Luciferase activity was normalized according to the mRNA levels of a *luciferase* gene. Red and blue rectangles represent wild-type (WT) CPEs, which are highlighted in Supplementary Fig. 3G, and an adenine mutated CPE#2 (mCPE#2), respectively. \*\**p* < 0.01; n.s., not significant (*n* = 3, mean ± SD, ANOVA followed by Tukey’s test)

The CPEB family as RNA-binding proteins are essential regulators of post-transcriptional gene expression, with functions including the polyadenylation of 3′UTRs and ribosome recruitment onto mRNA^13^. CPEB proteins recognize a pentanucleotide RNA sequence (5′-UUUUA-3′) in the 3′UTR of a target gene^14^. Five CPEB recognition motifs (CPEs) are found in the *NRF1*-3′UTR (numbers highlighted in red in Supplementary Fig. 3G). To identify the functional CPE, we initially checked by an RNA immunoprecipitation (RIP) assay that *NRF1* mRNA was precipitated with transiently expressed 3×Flag-CPEB3 proteins (Fig. 3G). We next confirmed that *NRF3* knockdown induced the translation *via NRF1*-3′UTR (Fig. 3H) using a translational assay with a luciferase reporter and also obtained results consistent with those of the *CPEB3* knockdown (Fig. 3I). We finally performed deletion and mutation analysis using this translational assay (Fig. 3J). The deletion of the region containing CPE#2–#5, but not deletion of the region containing CPE#3–#5, induced translation *via NRF1*-3′UTR in comparison with the full length (WT-Full vs. WT-Δ#2–#5 or WT-Full vs. WT-Δ#3–#5). The adenine mutation of the CPE#2 (UUUUA->AAAAA) also induced the translation *via* the full length, and #3–#5 deleted *NRF1*-3′UTR (WT-Full vs. mCPE#2-Full, or WT-Δ#3–#5 vs. mCPE#2-Δ#3–#5). These results clearly demonstrate that CPEB3 and CPE#2 in *NRF1*-3′UTR function as the *trans* regulator and its *cis*-element that are involved in NRF3-mediated repression of NRF1 translation.

### CPEB3 is a key factor for the maintenance of a basal proteasome activity and the poor prognosis of colorectal cancer patients

We investigated the impact of *CPEB3* overexpression on the basal expression of proteasome-related genes, the basal proteasome activity, and BTZ resistance. The basal mRNA levels of *PSMB3, PSMB7, PSMC2, PSMG2*, and *POMP* were not changed under a single knockdown of *NRF3* (Fig. 4A, Emp + siCont vs. Emp + siNRF3). Meanwhile, *CPEB3* overexpression significantly reduced these mRNA levels under a single knockdown of *NRF3* (Fig. 4A, CPEB3 + siCont vs. CPEB3 + siNRF3). Furthermore, *CPEB3* overexpression under *NRF3* single knockdown impaired the basal proteasome activity and the cancer cell resistance to BTZ (Fig. 4B and C). Finally, we validated the clinical relevance of the current findings in colorectal cancer where both *NRF* genes are highly expressed. Using the datasets archived at The Cancer Genome Atlas, we confirmed that colorectal cancer patients with higher *CPEB3*/*NRF3*-expressing tumors exhibit shorter overall survival rates, but higher *CPEB3*/*NRF1* expression is not associated with this prognosis (Fig. 4D and Supplementary Fig. 4A). These results clearly show that CPEB3 acts as a key factor for not only the complemental maintenance of basal proteasome activity in cancer cells by NRF1 and NRF3, but also the poor prognosis of colorectal cancer patients with high expression levels of *NRF3*, but not *NRF1*.

**Figure 4.**
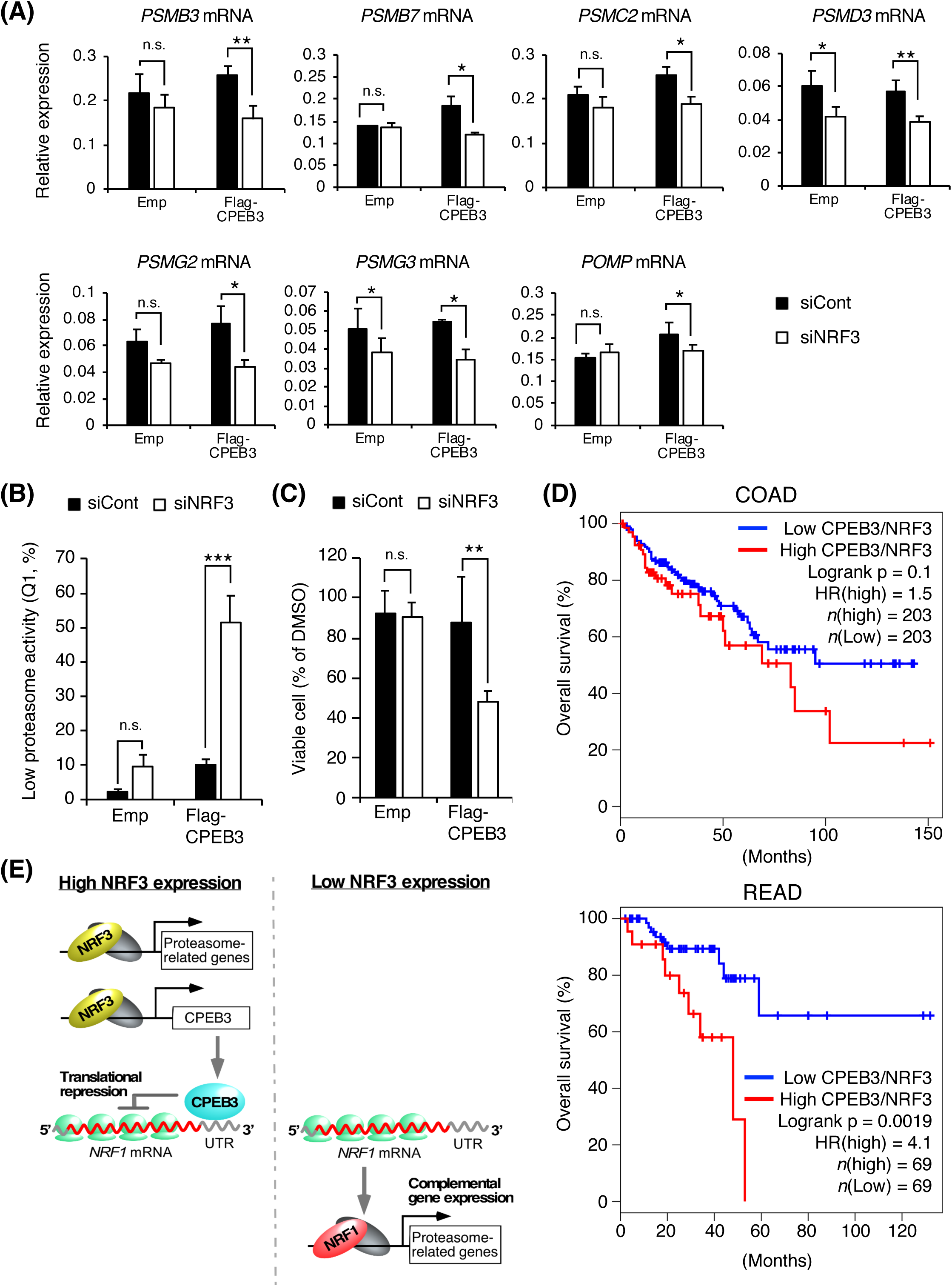
NRF3 and CPEB3 contribute to maintain proteasome activity in cancer cells and the prognosis of colorectal cancer patients. (**A–C**) Impact of *CPEB3* overexpression under *NRF3* knockdown on the expression of the proteasome-related genes (**A**), a proteasome activity (**B**), and the BTZ resistance (**C**). HCT116 or ZsPS cells were used in (**A** and **C**) or in (**B**), respectively. At 24 hours after NRF3 siRNA transfection, the cells were additionally transfected with the 3×Flag-CPEB3 plasmid and cultured for 24 hours. Control siRNA (siCont) and empty vector (Emp) were used as controls. In (**A**), the cells were subject to RT-qPCR using the primers for seven proteasome-related genes analyzed in Fig. 1C. mRNA levels of each proteasome subunit were normalized according to levels of *β-actin* mRNA. In (**B**), the fluorescence intensity derived from a ZsProSensor-1 reporter was measured using flow cytometry. The cell populations in Q1 enclosed by a red line are those with “low proteasome activity.” In (**C**), the cells were treated with 10 nM BTZ and further incubated for 24 hours. Then, the cells were subjected to cell viability assay using trypan blue staining. Cell viability was determined by the number of living cells with BTZ treatment normalized by that with DMSO. * *p* < 0.05, \*\**p* < 0.001, \*\*\**p* < 0.005; n.s., not significant (*n* = 3, mean ± SD, t-test). (**D**) Kaplan–Meier analysis comparing overall survival between higher and lower *CPEB3*/*NRF3* groups. The hazard ratio (HR) was calculated based on Cox’s proportional hazards model. Patients having colorectal adenocarcinoma (COAD) or rectal adenocarcinoma (READ) from The Cancer Genome Atlas. (**E**) Schematic model of the cross-talk mechanism between NRF1 and NRF3 to complementarily maintain a basal proteasome gene expression and activity in cancer cells through CPEB3-mediated translational repression.

## Discussion

Here, we demonstrated the complementary, but not simultaneous, regulatory mechanism of basal proteasome activity in cancer cells through the NRF3-CPEB3-NRF1 translational repression axis (Fig. 4E); In cancer cells, NRF3 induces the basal expression of proteasome-related genes in parallel with NRF1 translational repression by inducing *CPEB3* gene expression (left panel). If *NRF3* gene expression is reduced, NRF1 escapes from the CPEB3-mediated translational repression and complementarily plays a transcriptional role for the robust maintenance of basal proteasome activity in cancer cells (right panel). We also found that NRF1 induces *CPEB3* gene expression (Supplementary Fig. 3F) and that CPEB3 represses NRF1 translation (Fig. 3D and F). Although we have not confirmed yet whether NRF1 binds to the *CPEB3* promoter, we identified an NRF3-bound ARE in the *CPEB3* promoter (Fig. 3B). Considering that NRF1 and NRF3 bind to the ARE sequence, these results suggest the possibility of negative feedback regulation of NRF1 through CPEB3-mediated translational repression (Supplementary Fig. 4B).

*NRF1* is ubiquitously expressed in normal tissues, and mice in which it is knocked out suffer embryonic lethality^15^. Meanwhile, *NRF3* expression levels are low except in several tissues such as the placenta^8^, and *Nrf3* knockout mice do not exhibit any apparent abnormalities under normal physiological conditions^16–18^. However, *NRF3* is highly expressed in many types of cancer cells, implying that the proteasome in cancer or normal cells is maintained through the NRF3-CPEB3-NRF1 translational repression axis or the negative feedback regulation of NRF1. Indeed, we revealed a clinical association of higher *CPEB3*/*NRF3* expression, but not higher *CPEB3*/*NRF1* expression, with poor prognosis of cancer patients (Fig. 4D and Supplementary Fig. 4A). Therefore, our findings suggest the possibility that while NRF1 maintains a basal proteasome activity within an appropriate range for normal development, NRF3 adjusts the proteasome in the quality and quantity for cancer development.

We guess another biological meaning of our findings in cancer. The functional difference between NRF3 and NRF1 is not in the regulation of the proteasome activity rather in the expression of cancer-related genes such as *VEGFA* and *DUX4* that other groups have recently identified as NRF3-specific target genes^19,20^. Meanwhile, cancer cells have to keep NRF1 as a backup for proteasome regulation, because the proteasome activity is crucial for more rapid proliferation of cancer cells^21^. Thus, the NRF3-CPEB3-NRF1 translational repression axis enables cancer cells to grow more rapidly through NRF3-mediated signals and to maintain the proteasome activity through NRF1 immediately in case NRF3 does not work. Further study is needed to confirm this point.

Unexpectedly, *NRF1* knockdown did not affect NRF3 protein levels (Fig. 2A, Supplementary Fig. 2A), although *NRF1* knockdown reduced *CPEB3* mRNA levels (Supplementary Fig. 3F) and *NRF3* gene possessed several CPEs in its 3′UTR (blue highlighted numbers in Supplementary Fig. 3G). CPEB3 belongs to the CPEB family consisting of four paralogs, CPEB1, CPEB2, CPEB3 and CPEB4^22^. According to the binding RNA sequence, the CPEB family is functionally categorized into two groups, CPEB1 and CPEB2-4^23^, implying that CPEB2 and/or CPEB4 compensates for the reduction of the *CPEB3* gene by NRF1 depletion. An alternative possibility is a conversion of CPEB function in a cellular context-dependent manner^13^. In maturing mouse and *Xenopus* oocytes, CPEB acts as the translational activator through phosphorylation by aurora kinase^24,25^. Therefore, there is an unknown mechanism of cross-talking between NRF and CPEB protein families.

## Materials and Methods

### Cell lines

HCT116 (human colorectal carcinoma), T98G (human glioblastoma multiforme), and MCF-7 (human breast cancer) cells were cultured in DMEM/high glucose (Wako) supplemented with 10% FBS (Nichirei), 40 µg/ml streptomycin, and 40 units/ml penicillin (Life Technology). H1299 (human non-small cell lung cancer) cells were cultured in RPMI-1640 (Nacalai Tesque) supplemented with 10% FBS (Nichirei), 40 µg/ml streptomycin, and 40 units/ml penicillin (Life Technology). To generate the cells stably expressing a ZsProSensor reporter (ZsPS cells), HCT116 cells were transfected with the proteasome sensor vector (Clontech). To generate *NRF3*- and *GFP*-overexpression cells, H1299 cells were transfected with the p3×FLAG-CMV 10 vector (Sigma-Aldrich) containing human full-length NRF3 or GFP. The transfected cells were selected with G-418.

### Plasmid construction and mutagenesis

The 3×Flag-CPEB3 plasmid was generated by subcloning the PCR-amplified human CPEB3 cDNA into the p3×FLAG-CMV 10 vector (Sigma-Aldrich). The human CPEB3 cDNA was synthesized using the indicated primers. Forward: 5′-TTTGAATTCAATGCAGGATGATTTAC-3′, Reverse: 5′-AAAGGATCCTCAGCTCCAGCGGAAC-3′. The luciferase reporter driven by the *NRF1*-3′UTR-Full was generated by PCR amplification of human genomic DNA using the indicated primers (Forward: 5′-GGCCGGCCGCCTGGGAAGAAGGGGGTT-3′, Reverse: 5′-GGCCGGCCTTTTTTTTTTTTTTTTTTACAATGAGTCCA-3′), and cloned into pGL3-Control Vector (Promega). The deletion of *NRF1*-3′UTR (*NRF1*-3′UTR-Δ#2-#5 or Δ#3-#5) was performed by a PCR-based method using the indicated primers. Forward: 5′-TTTTTGGCCGGCCCCTGGGGAAGAAG-3′, Reverse: 5′-TTTTTGGCCGGCCGACATTGTAGTCC-3′ (*NRF1*-3′UTR-Δ #2-#5), or 5′-TTTTTGGCCGGCCGCACTCCCAGGTT-3′ (*NRF1*-3′UTR-Δ#3-#5). The adenine mutation of CPE#2 in *NRF1*-3′UTR were introduced by site-directed mutagenesis using PCR. The primer sequence was as follows (altered nucleotides are underlined): 5′-ACAATGTCTTTAT<u>AAAA</u>AACTGTTTGCCA-3′. All constructs were confirmed by sequencing.

### Antibody

The antibodies utilized in the current immunoblot analysis were anti-α-tubulin (DM1A; Sigma-Aldrich), anti-NRF1 (D5B10; Cell Signaling Technology), anti-NRF3 (#9408), and anti-FLAG (M2; Sigma-Aldrich). A monoclonal NRF3 antibody (#9408) raised against human NRF3 (amino acids 364-415) was generated as described previously^26^.

### Transfection

The transfection of the plasmid DNA and short interfering RNA (siRNA) was performed using polyethylenimine (PEI) and RNAiMAX (Invitrogen), respectively. The sequences of the siRNA duplexes are listed in Supplementary Table 3.

### RNA extraction and real-time quantitative PCR (RT-qPCR)

Total RNA was extracted and purified using ISOGEN II (NIPPON GENE) according to the manufacturer’s instructions. Aliquots of total RNA (1 μg) were reverse transcribed using pd(N)6 random primer (Takara Bio) and Moloney murine leukemia virus (M-MLV) reverse transcriptase (Invitrogen) with 250 μM deoxy nucleoside triphosphate (dNTP, Takara Bio) concentration, according to the manufacturer’s instructions. RT-qPCR was performed with SYBR Premix Ex Taq II (Takara Bio) and primers for genes using a Thermal Cycler Dice Real Time System (Takara Bio). Each gene expression level in human cells was normalized to the mRNA levels of human *β-actin* gene. qPCR primer sequences are described at Supplementary Table 3.

### Immunoblot analysis

To prepare whole cell extracts, the cells were lysed with SDS sample buffer (50 mM Tris-HCl [pH 6.8], 10% glycerol and 1% SDS). The protein quantities in cell extracts were measured with a BCA kit (ThermoFisher Scientific). The proteins were separated by sodium dodecyl sulfate-polyacrylamide gel electrophoresis (SDS-PAGE) and transferred to PVDF membranes (Immobilon-P transfer membrane, EMD Millipore corporation). After blocking the membranes with Blocking One (Nacalai Tesque) at 4°C for overnight, the membranes were incubated with a primary antibody, washed with TBS-T (20 mM Tris-HCl [pH 7.6], 137 mM NaCl, 0.1% Tween20) and were incubated with a horseradish peroxidase-conjugated secondary antibody (Invitrogen). The blots were washed with TBS-T and developed with enhanced chemiluminescence (GE Healthcare).

### Cycloheximide chase experiments

HCT116 cells were transfected with the indicated siRNA. At 48 hours after transfection, the cells were treated with 20 µg/ml cycloheximide (CHX), and the whole cell extracts were prepared at the indicated time points. The immunoblot analysis was conducted with the indicated antibodies.

### Protein synthesis assays

The protein synthesis assay was conducted using Click-iT Plus OPP Protein Synthesis Assay Kit (Invitrogen) according to the manufacturer’s protocol. HCT116 cells were transfected with the indicated siRNA. At 48 hours after transfection, the cells were treated with O-propargyl-puromycin (OPP; 20 µM) for 1 hour at 37°C and washed twice with PBS. The cells were fixed by using 4% paraformaldehyde for 15 min at room temperature, followed by washing them twice with PBS (containing 0.5% BSA) and then permeabilizing with PBSTx (0.5% TritonX-100/PBS) for 15 min at room temperature. The cells were washed twice with PBS (containing 0.5% BSA). After their treatment with Click-iT Reaction Mixtures for 30 min at room temperature in a dark place, the cells were then washed with Click-iT Reaction Rise Buffer. Finally, the cells were washed with PBS and subjected to the protein synthesis assay using BD FACS Aria-II (Becton Dickinson).

### Polysome fractionation assays

HCT116 cells were treated with 100 µg/ml cycloheximide (CHX) in culture medium for 5 min at 37°C. The cells were washed twice with PBS containing CHX (100 µg/ml) and lysed with lysis buffer (15 mM Tris-HCl [pH 7.5], 300 mM NaCl, 25 mM EDTA, 1% Triton X-100, 0.5 mg/ml heparin, 100 µg/ml CHX) with 1× EDTA-free protease inhibitor cocktail (Nacalai Tesque). The lysate was centrifuged at 9,300 × *g* for 10 min at 4°C and loaded onto a linear 20–50% sucrose gradient buffer in 15 mM Tris-HCl [pH 8.0], 25 mM EDTA and 300 mM NaCl. Centrifugation was conducted at 40,000 rpm for 2 hours at 4°C in a SW-41 Ti rotor (Beckman Coulter), and fractions were collected from the top of the gradient (#1 to #20). The RNA from each fraction was subjected to RT-qPCR analysis. The quantified mRNA was normalized by the input. qPCR primer sequences are described at Supplementary Table 3.

### DNA microarray analysis

Total RNA was processed with the Ambion WT Expression Kit (Affymetrix) according to the manufacturer’s instructions. cRNA was then fragmented, labelled, and hybridized to the Affymetrix Human Gene 1.0 ST Arrays using the Gene Chip WT Terminal Labeling and Hybridization Kit (Affymetrix). GeneChip fluidics station 450 was used for processing of the arrays and fluorescent signals were detected with the GeneChip scanner 3000-7 G. Images were analyzed with the GeneChip operating software (Affymetrix).

Expression console and Transcription analysis console (Affymetrix) were used to analyze the data. PANTHER Classification System with the theme “Molecular function” was used for gene ontology analysis of the genes whose expression was reduced upon siRNA-mediated *NRF3*-knockdown HCT116 cells (fold change ≧ 1.4) and increased by *NRF3*-stable overexpression H1299 cells (fold change ≧ 1.4). The expression data of all genes in siRNA-mediated *NRF3*- or *NRF1*-knockdown HCT116 cells were subjected to GSEA using open source software v3.0^27^ and the gene set containing 42 proteasome-related genes in Supplementary Table 1.

### Chromatin immunoprecipitation (ChIP) qPCR

The cells were fixed with 1% formaldehyde for 10 min at room temperature, and then glycine was added to a final concentration of 0.125 M. The cells were lysed with cell lysis buffer (5 mM Tris–HCl [pH 8.0], 85 mM KCl, 0.5% NP-40) with protease inhibitors (Nacalai Tesque), and then centrifuged at 2,000 rpm at 4°C for 3 min. The pellets were further lysed with nuclei lysis buffer (50 mM Tris–HCl [pH 8.0], 10 mM EDTA, 1% SDS) with protease inhibitors (Nacalai Tesque), and the lysates were sonicated. After centrifugation at 15,000 rpm at 8°C for 10 min, the supernatants were collected. The supernatants were diluted in ChIP dilution buffer (16.7 mM Tris–HCl [pH 8.0], 167 mM NaCl, 1.2 mM EDTA, 1.1% TritonX-100, 0.01% SDS). The diluted samples were precleared with 20 µl of Dynabeads Protein G (ThermoFisher Scientific), and then the supernatants (used as an input sample) were incubated with 2 µg of anti-NRF3 antibody. The immunocomplexes were collected by incubation with 20 µl of Dynabeads Protein G (ThermoFisher Scientific), and then washed with the following buffers: low salt wash buffer (20 mM Tris–HCl [pH 8.0], 150 mM NaCl, 2 mM EDTA, 1% Triton X-100, 0.1% SDS), high salt wash buffer (20 mM Tris–HCl [pH 8.0], 500 mM NaCl, 2 mM EDTA, 1% Triton X-100, 0.1% SDS), and LiCl wash buffer (10 mM Tris–HCl [pH 8.0], 250 mM LiCl, 1 mM EDTA, 1% sodium deoxycholate, 1% NP-40). Finally, the beads were washed twice with 1 ml of TE buffer (10 mM Tris–HCl [pH 8.0], 1 mM EDTA). The immunocomplexes were then eluted by adding 200 µl of elution buffer (50 mM NaHCO_3_, 1% SDS). After the reverse cross-linking by adding 200 mM NaCl, the remaining proteins were digested by adding proteinase K. For quantification of NRF3-binding to the target regions, RT-qPCR was performed using the purified DNA with the primers described at Supplementary Table 3.

For ChIP-sequencing, the libraries were prepared from 500 pg of immunoprecipitated DNA fragments using the KAPA Hyper Prep Kit (KAPA Biosystems). The libraries were applied to Single-end sequencing for 93 cycles on HiSeq2500 (Illumina). All sequence reads were extracted in FASTQ format using BCL2FASTQ Conversion Software 1.8.4 in the CASAVA 1.8.2 pipeline. Mapping was performed by BWA (version 0.5.9rc1) using the reference human genome, NCBI build 37 (hg19), and peak call was conducted using MACS (version 1.4.2).

### RNA immunoprecipitation (RIP)

HCT116 cells were transfected with empty vector or 3×Flag-CPEB3 plasmid. At 48 hours after transfection, cells were lysed in cell lysis buffer containing protease and RNase inhibitors. Lysates were sonicated for 10 min at low intensity and then precleared. Samples were immunoprecipitated using anti-FLAG antibody coupled to Dynabeads Protein A (Invitrogen) for 3 hours at 4°C. RNA bound the FLAG-beads was purified by a phenol/chloroform extraction, and was subjected to RT-qPCR analysis. qPCR primer sequences are described at Supplementary Table 3.

### Luciferase reporter assays

Cells expressing the reporters indicated in the legends for Fig. 3H-J were lysed, and the luciferase activities were measured with the PicaGene luciferase assay system (Toyo Ink) and a microplate reader (Synergy HTX, Bio Tek Instruments).

### Proteasome activity analysis using a ZsProSensor reporter

The ZsProSensor-1 protein is a fusion of the green fluorescent protein ZsGreen and mouse ornithine decarboxylase (MODC), and can be degraded by the proteasome without being ubiquitinated. ZsPS cells were transfected with the indicated siRNA or plasmid. At 48 hours after transfection, the cells were collected, followed by washing them twice with STM (2% FBS in cold PBS). 500 μl of STM containing 2 μg/ml of propidium idodide (PI) was added to each sample. After their treatment at 4°C in a dark place, samples were subjected to the proteasome activity analysis using BD FACS Aria-II (Becton Dickinson).

### *in vitro* proteasome activity assays

*in vitro* proteasome activity assays were based on a glycerol density gradient centrifugation and fluorogenic peptidase assays as described previously^28^. After centrifugation at 26,000 rpm for 22 hours in a Beckman SW40Ti swing rotor, the gradient was manually separated into 20 fractions of 500 µl each. Since the 26S proteasome, which is made up of a 20S proteasome and one or two 19S-RPs, is contained in fraction #13 and #14, the average of the fluorescent intensity was calculated as the proteasome activity as follows: 30 µl of each fraction sample was transferred to a 96-well BD Falcon microtiter plate (BD Biosciences), and mixed with 2 mM ATP and 0.1 mM fluorogenic peptide substrate Suc-Leu-Leu-Val-Tyr-AMC (Peptide Institute). Fluorescence (380 nm excitation, 460 nm emission) was monitored on a microplate fluorometer (Synergy HTX, BioTek Instruments) every 5 min for 1 hour^29^.

### Cell viability assays using Trypan Blue staining

HCT116 cells were transfected with siRNA and/or plasmid DNA under the conditions indicated in the legends for Figs. 1B and 4C. After the treatment with 5 nM or 10 nM bortezomib (BTZ, Peptide Institute), the cells were stained by Trypan Blue and automatically counted using a TC20 Automated Cell Counter (Bio-Rad Laboratories).

### Statistics and human cancer database

The unpaired Student’s t-test was used to compare the two groups, and one-way analysis of variance followed by Tukey’s post-hoc test was used to compare multiple groups. The GEPIA, a web server for cancer and normal gene expression profiling and interactive analyses^10^ was used for Kaplan–Meier analyses.

## Acknowledgments

We thank Hiderori Watanabe, Mika Matsumoto and Junko Naritomi for experimental supports. This work was supported in part by grant-in-aid for Young Scientists (B) (17K18234 to T.W.); grant-in-aid for Scientific Research (C) (19K07650 to T.W.); The Uehara Memorial Foundation (to T.W.); grant-in-aid for Scientific Research (B) (16H03265 to A.K.); grant-in-aid for Challenging Research (Exploratory) (19K22826 to A.K.); grant-in-aid for JSPS Fellows (18J20672 to A.H.).

## Contributions

T.W., H.K., and Y.T. conceived the project; T.W. and H.K. designed experiments; T.W., H.K., M.H., A.H., N.N., Y.T., N.T. and A.W. performed experiments and analyzed data; T.W. and A.K. wrote the manuscript; T.W. and A.K. provided funding; A.K. provided supervision.

---

**Supplementary Figure 1.** NRF1 and NRF3 function on the basal gene induction of proteasome-related factors. (**A**) mRNA levels of the ZsProSensor reporter gene. Parental and ZsPS cells were analyzed using RT-qPCR. mRNA levels of the reporter gene were normalized according to levels of *β-actin* mRNA. \*\*\**p* < 0.005 (*n* = 3, mean ± SD, t-test) (**B**) The *in vitro* activity of the 26S proteasome fractionated from *NRF* knockdown cells. At 24 hours after indicated siRNA transfection into HCT116 cells, the cell extracts were manually fractionated into 20 fractions using a 10%–40% glycerol gradient centrifugation and the average of the fluorescent intensity in fraction #13 and #14 was calculated as the 26S proteasome activity. siNRF1/3 is a double knockdown of *NRF1* and *NRF3*. Control siRNA (siCont) was used as a control. \**p* < 0.05; n.s., not significant (*n* = 3, mean ± SD, t-test) (**C**) Impact of MG-132 treatment of proteasome activity in ZsPS cells. At 24 hours after treatment with 1 μM MG-132, the cells were analyzed using flow cytometry. DMSO was used as control. The cell populations in Q1 enclosed by a red line are shown as “low proteasome activity.” \*\*\**p* < 0.005 (*n* = 3, mean ± SD, t-test) (**D** and **E**) Enrichment plots for the gene sets involved in a proteasome. DNA microarray data using *NRF1*- or *NRF3*-knockdown HCT116 cells are shown in (**C**) and (**D**), respectively. Control siRNA (siCont) was used as a control.

**Supplementary Figure 2.** NRF3 does not change polysome formation or global translation as well as protein degradation of NRF1. (**A** and **B**) Impact of *NRF* knockdown on protein and mRNA levels of NRF1 or NRF3 in T98G (glioblastoma multiforme) and MCF-7 (breast cancer) cells. At 48 hours after siRNA transfection, cells were analyzed by immunoblotting (**A**) and RT-qPCR (**B**). Control siRNA (siCont) was used as a control. mRNA levels of each NRF were normalized using *β-actin* mRNA. n.s., not significant (*n* = 3, mean ± SD, ANOVA followed by Tukey’s test in **B**) (**C** and **D**) Impact of *NRF3* overexpression on protein and mRNA levels of NRF1. *NRF3*-overexpression H1299 cells were analyzed by immunoblotting (**C**) and RT-qPCR (**D**). *GFP*-overexpression H1299 cells were used as controls. mRNA levels of NRF1 were normalized by *β-actin* mRNA. n.s., not significant (*n* = 3, mean ± SD, t-test in **D**) (**E**) Distribution of ribosomal RNAs in HCT116 cells transfected with NRF1 or NRF3 siRNA. At 48 hours after siRNA transfection, cells were analyzed using sucrose gradient centrifugation followed by agarose gel electrophoresis. Control siRNA (siCont) was used as a control. (**F**) Impact of *NRF* knockdown on a global translation. At 48 hours after siRNA transfection, the cells were stained using an OPP-based protein synthesis assay followed by flow cytometry. Control siRNA (siCont) and CHX were used as controls. n.s., not significant (*n* = 3, mean ± SD, ANOVA followed by Tukey’s test).

**Supplementary Figure 3.** The translational regulator CPEB3 does not affect global protein synthesis. (**A**) Venn diagram combining two independent DNA microarray datasets of *NRF3*-knockdown (NRF3KD) HCT116, and *NRF3*-overexpression (NRF3OE) H1299 cells. *GFP*-overexpression H1299 cells were used as a control. (**B**) Gene ontology analysis of 146 common genes using the PANTHER Classification System. The term “Molecular function” was selected. (**C**) Knockdown efficiency of CPEB3 siRNAs. At 48 hours after siRNA transfection, cells were analyzed using RT-qPCR. mRNA levels of *CPEB3* were normalized according to levels of *β-actin* mRNA. Control siRNA (siCont) was used as a control. \**p* < 0.05 (*n* = 3, mean ± SD, ANOVA followed by Tukey’s test). (**D**) Distribution of ribosomal RNAs in HCT116 cells transfected with CPEB3 siRNA. At 48 hours after siRNA transfection, cells were analyzed using sucrose gradient centrifugation followed by agarose gel electrophoresis. Control siRNA (siCont) was used as a control. (**E**) Impact of *CPEB3* knockdown on a global protein synthesis. At 48 hours after siRNA transfection, the cells were stained using an OPP-based protein synthesis assay and analyzed by flow cytometry. Control siRNA (siCont) and CHX were used as controls. n.s., not significant (*n* = 3, mean ± SD, ANOVA followed by Tukey’s test). (**F**) Impact of *NRF1* knockdown on mRNA levels of *CPEB3* in HCT116 cells. At 48 hours after siRNA transfection, the cells were analyzed by RT-qPCR. Control siRNA (siCont) was used as a control. mRNA levels of *CPEB3* were normalized by *β-actin* mRNA. \**p* < 0.05 (*n* = 3, mean ± SD, t-test) (**G**) DNA sequence alignment of *NRF1*- and *NRF3*-3′UTR. CPEB recognition motifs (5′-UUUUA-3′) in *NRF1*- and *NRF3*-3′UTR were highlighted by red and blue, respectively.

**Supplementary Figure 4.** Clinical association of *NRF1* and *CPEB3* expression levels with the prognosis of colorectal cancer patients. (**A**) Kaplan–Meier analysis comparing overall survival between higher and lower *CPEB3*/*NRF1* groups. The hazard ratio (HR) was calculated based on Cox’s proportional hazards model. Patients having colorectal adenocarcinoma (COAD) or rectal adenocarcinoma (READ) from The Cancer Genome Atlas. (**B**) Schematic model of the negative feedback regulation of NRF1 through CPEB3-mediated translation repression.

